# Long-term dynamics of the “*Serratia marcescens* complex” in the hospital-built environment

**DOI:** 10.1101/2023.10.15.562376

**Authors:** Sonia Aracil-Gisbert, Miguel D. Fernández-De-Bobadilla, Natalia Guerra-Pinto, Silvia Serrano-Calleja, Ana Elena Pérez-Cobas, Cruz Soriano, Raúl de Pablo, Val F. Lanza, Blanca Pérez-Viso, Sandra Reuters, Henrik Hasman, Rafael Cantón, Fernando Baquero, Teresa M. Coque

## Abstract

*Serratia marcescens* is an opportunistic pathogen historically associated with abrupt, unpredictable, and severe outbreaks in hospital intensive care units (ICUs) and, more recently, with the spread of acquired genes encoding carbapenem resistance. However, the population biology and ecology of *S. marcescens* in the hospital ecosystem is still poorly understood. Here, we combine epidemiological information of 1417 *Serratia* isolates collected from the sinks of a large ICU ward that underwent significant demographic and operational changes (2019-2020) and 99 non-redundant outbreak/non-outbreak isolates from the same hospital (2003-2019), with genomic data of 165 isolates. We first report hospital sinks as reservoirs of heterogeneous and coexistent populations of the *S. marcescens* complex (SMC). A novel SMC clade congruent with the recently described *Serratia nevei* species is predominant, exhibiting chromosomal AmpC β-lactamase with an unusual basal expression in contrast to one of the major features of *S. marcescens*. Persistent *Serratia* sink strains are identical to those involved in clonal and polyclonal outbreaks of VIM-1 and OXA-48 producers since at least 2017. The “source-sink” dynamics of SMC populations able to acquire the highly conserved plasmids such as IncL carrying *bla*_VIM-1_ or *bla*_OXA-48_ offer novel insights that might improve interventions to control outbreaks and treat Serratia infections in high-risk hospital areas.

---

The *Serratia* genus belongs to the order *Enterobacterales*^1^, comprising species widely distributed in water, soil, and the rhizosphere, which can become opportunistic pathogens for animals, humans, and plants^2–4^. In the health sector, *Serratia* accounts for only 1%–2% of hospital-acquired infections, which are often associated with unpredictable outbreaks of high morbidity and mortality in pediatric or intensive care unit (ICU) wards. Contamination from medical devices (e.g. nebulizers, catheters, fomites) and via healthcare workers are well-known transmission pathways^3^. Hospital water sinks are increasingly recognized as reservoirs of multidrug-resistant *Serratia* and other *Enterobacterales* during outbreaks^5–7^. Nonetheless, the dynamics of *Serratia* in the hospital environment remain under-studied.

According to the National Center for Biotechnology Information (NCBI) Taxonomy databases, the *Serratia* genus comprises 26 species, with *Serratia marcescens* being by far the predominant cause of human infections^3^. An increasing number of genomic studies have led to the description of several novel species and the reclassification of others within the *Serratia* genus in recent years^8^. Phylogenetic analysis have lately categorized several emblematic *Serratia* species within a “*Serratia marcescens* complex” (SMC), thus highlighting the remarkable diversification of *S. marcescens* associated with its adaptation to various environments^2,9,10^.

One of the main features of *S. marcescens* and other *Enterobacterales*, known as the ESCPM group (*Enterobacter* spp., *S. marcescens, Citrobacter freundii, Providencia* spp., *Morganella morganii*), is their intrinsic resistance to β-lactams (amino- and carboxy-penicillins, amoxicillin-clavulanate [AMC], first- and second-generation cephalosporins [1GC, 2GC]), due to the production of a chromosomally encoded inducible AmpC β-lactamase^11,12^. The induction of AmpC β-lactamase is of outstanding clinical relevance due to the risk of AmpC derepression upon therapy^13^. Such risk, and the increasing occurrence during the 1990s of plasmid-mediated extended-spectrum β-lactamases (ESBLs) conferring resistance to third-generation cephalosporins (3GCs), led to the discouraged use of cephalosporins and the increased use of carbapenems for *Serratia* infections^14^. Consequently, the emergence and dramatic increase of carbapenem-resistant strains causing severe nosocomial infections has increasingly been reported^15^.

In this work, we analyze the diversity of *Serratia* isolated in the sinks of the largest ICU ward in our hospital, where recurrent outbreaks of ESBL and carbapenemase producers occurred over time^16–19^. The goals of our study were to i) evaluate the role of ICU-built environment as a “source” and “sink” of *Serratia* in the hospital environment, and ii) evaluate the role of *Serratia* as a reservoir of clinically relevant plasmids encoding ESBL/carbapenemase genes.

## RESULTS

### Spatiotemporal distribution of *Serratia* spp. in the intensive care unit

The 1460 *Serratia* isolates in this study were recovered during a 34-month prospective study in the largest ICU ward of a tertiary hospital (14 rooms, 2417 samples of “dry” and “wet” abiotic surfaces, 13180 bacterial isolates), starting shortly after an outbreak caused by *S. marcescens* producing VIM-1^16^. Several demographic changes in this ward reflect the 3 study periods. Periods A (March 2019-February 2020) and B (July 2020-February 2021) cover the ICU occupancy before and during the SARS-CoV-2 epidemic, respectively. Afterward, the ICU was emptied, and patients were transferred to another hospital floor with operational transformation of the ward (period C, March December 2021). This offers a unique chance to identify high-risk hospital environment reservoirs of opportunistic pathogens.

*Serratia* was mainly recovered from “wet” surfaces (448/1126 samples, 39.8%; 1417 isolates), namely sink drains (253/448, 56.5%), washbasin surfaces (168/448, 37.5%), and p-trap samples (27/448, 6.0%). Detection on “dry” surfaces was sporadic (19/1291, 1.5%; 43 isolates). Then, we focused on analyzing *Serratia* from sink samples. **Fig. 1** shows the spatiotemporal distribution of these isolates. The highest recovery rate of *Serratia* occurred during period A, overlapping with the ICU outbreak by *S. marcescens* producing VIM-1 (385/746, 51.6%; 14/14 sinks). During periods B and C, the isolation frequency drastically decreased (58/247, 23.5%, 13/14 sinks during period B; 5/133, 3.8%, 1/14 sinks during period C). Most of the 1417 colonies tested were preliminarily identified by matrix-assisted laser desorption/ionization-time of flight (MALDI-TOF) mass spectrometry (MS) as *S. marcescens* (n=1261, 89.0%), followed by *Serratia ureilytica* (n=151, 10.7%), *Serratia nematodiphila* (n=4, 0.3%), and *Serratia entomophila* (n=1, 0.1%). *S. marcescens* and *S. ureilytica* isolates were found in all sinks, whereas *S. entomophila* was detected only once, and *S. nematodiphila* was isolated in a single room during period C.

**Fig 1.**
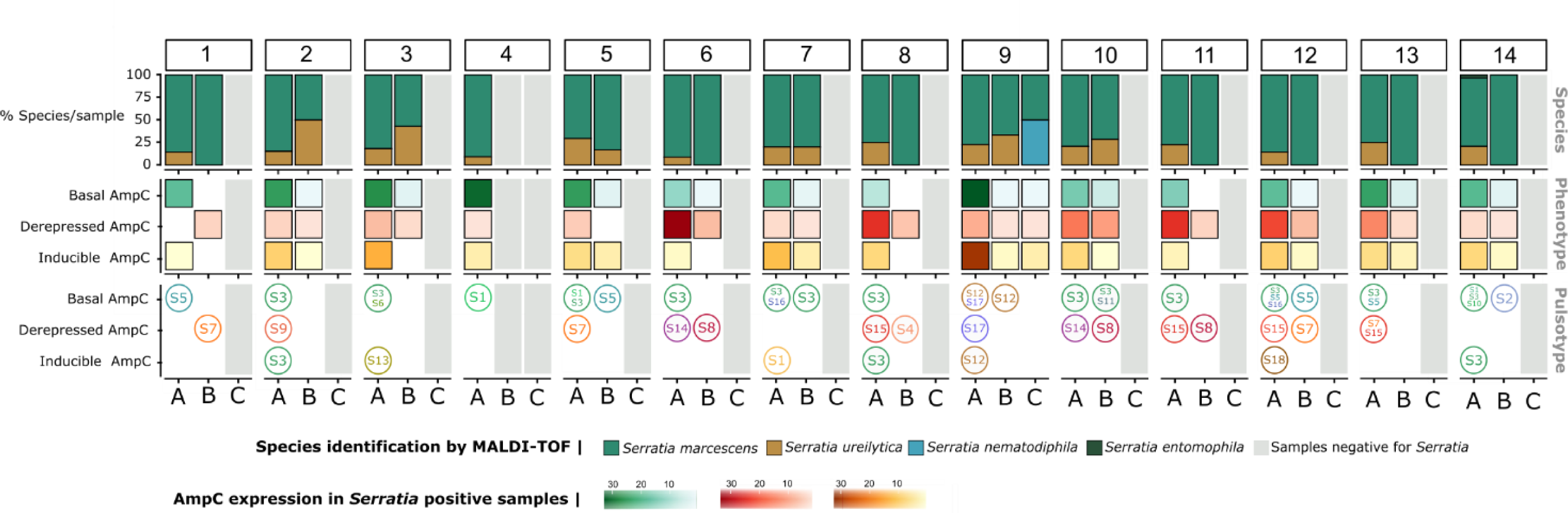
Spatiotemporal distribution of *Serratia* in sinks at Hospital Ramón y Cajal. Columns labeled by numbers embedded in boxes represent the 14 intensive care unit (ICU) rooms. Capital letters designate the study periods: A (March 2019-February 2020), B (July 2020-February 2021), and C (March 2021-December 2021), according to patient occupancy and use of the ICU hospital space, namely, ICU occupancy before (A) and after SARS-CoV lockdown (B), and patient clearing and conversion of the ward for other purposes unrelated to inpatient care (C). The first line shows the diversity and percentage of each *Serratia* species according to matrix-assisted laser desorption/ionization time of flight (MALDI-TOF) mass spectrometry. The second row reflects the diversity of AmpC phenotypes, each phenotype being represented by a color: i) basal expression of the AmpC enzyme (AMC^S^+FOX^S^+3GC^S^); ii) derepressed AmpC expression (AMC^R^+FOX^R^+3GC^R^+4GC^R^±carbapenems^R^), and iii) inducible AmpC β-lactamase expression (AMC^R^+FOX^R^+3GC^S^). The third row represents the most common XbaI digested genomic DNA patterns (pulsotypes, PT) of isolates with distinct AmpC phenotypes.

### *Serratia*’s β-lactam phenotype diversity

We identified 3 phenotypic patterns compatible with i) basal expression of the AmpC enzyme (susceptibility [S] to AMC, cefoxitin [FOX], and 3GC: AMC^S^+FOX^S^+3GC^S^); ii) inducible AmpC β-lactamase expression (resistance [R] to AMC and FOX and 3GC^S^: AMC^R^+FOX^R^+3GC^S^); and iii) derepressed AmpC expression (AMC^R^+FOX^R^+3GC^R^±4GC^R^±carbapenems^R^) due to the presence of the carbapenemase VIM-1). The first phenotype is similar to that of *Escherichia coli* but is exceptional for the ESCPM group and has not been reported for *S. marcescens*^12^. Isolates with AmpC basal expression exhibit minimum inhibitory concentrations to aminopenicillins lower than those producing inducible AmpC or hyperproducing the enzyme (2-8 mg/mL vs. 16->64 mg/mL, respectively).

All AmpC expression patterns were detected throughout the study, although isolates with the basal phenotype predominated (286/448 sink samples, 63.8%) over the “classical” AmpC phenotypes, either inducible (144/448, 32.1%) or derepressed (180/448, 40.2%). It is of note that *Serratia* isolates exhibiting all 3 AmpC phenotypes were regularly detected in the same sink sample, although the frequency of each phenotype varied among sinks. A progressive decrease in the recovery of *Serratia* inducible phenotypes as well as an increase of derepressed AmpC isolates during period B was observed (**Fig. 1**). Resistance to chloramphenicol, gentamicin, and sulfamethoxazole-trimethoprim was observed only among AmpC-derepressed *Serratia*, whereas susceptibility to ciprofloxacin and tetracycline varied significantly within and between groups.

### Genomic diversity of *Serratia* strains

We selected a sample of 94/1417 sink isolates for further analysis. The selection was randomly made among isolates representing the spatiotemporal distribution of species and phenotypes after eliminating redundancies (same antibiogram, morphology, and sampling site). Then, we preliminary analyzed the clonal relationship by pulsed-field gel electrophoresis (PFGE) typing and grouped the 94 isolates into 18 pulsotypes (PTs) of which 15 corresponded to *S. marcescens* and 3 to *S. ureilytica* strains according to MALDI-TOF. Some PTs, the predominant PT-S3 and PT-S1, PT-S5, PT-S7, PT-S14, and PT-S15, were detected in the majority of sinks during the whole study period while others (PT-S2, PT-S4, PT-S6, PT-S9, PT-S10, PT-S12, PT-S13, PT-S17, PT-S18) were persistently recovered in specific sinks (**Fig. 1**). A sample of 66 isolates representing the spatiotemporal distribution, the diversity of β-lactam-phenotypes, and the PFGE patterns were fully sequenced.

To further explore the phylogenetic diversity and understand how variation in habitat quality influences the growth or decline of *Serratia* populations in the hospital environments (“the source-sink dynamics” ecological model^20^), we comparatively analyzed the genomes of the 66 ICU-sink isolates with 99 clinical isolates representing the epidemiology of *Serratia* in our institution in the last years. They included 94 isolates causing independent episodes of bloodstream infection (BSI, 2003-2015) and 5 strains recovered during outbreaks of VIM-1 and OXA-48 producers (2016-2018)^16,19^. Most of the clinical isolates were collected in ICU wards. The dataset of the 165 sequenced isolates is summarized in **Table S1**.

The chromosome length of the genomes analyzed ranged from 4.9 to 5.4 Mb (59.4% G+C content). The core-based phylogenetic tree represented in **Fig. 2** shows 2 major clades arbitrarily designated as clade 1 and 2, each split into 2 subclades assigned the capital letters “A” and “B” (subclades 1A, 1B, 2A, 2B). We overlayed these clades/subclades with available phenotypic info and their similarity to available genomes in the GenBank databases. We found that the subclade 1A corresponded with isolates identified as *S. marcescens* (Sm), subclade 1B with *S. ureilytica* (Su), and subclades 2A and 2B with the recently described *Serratia nevei* (Sn) species. The pairwise ANI scores within and between subclades were >95%, indicating that all belong to the same species (**Fig. S1**). We arbitrarily called them “lineage Sm” (1A), “lineage Su” (1B), and “lineages Sn-like” (2A and 2B) to link the clades with known taxons and facilitate the understanding of the results in the context of the available knowledge. *Serratia* isolated from sinks were mostly grouped in “lineages Sn” (2A and 2B) with a few isolates distributed into “lineage Su” (1B).

**Fig 2.**
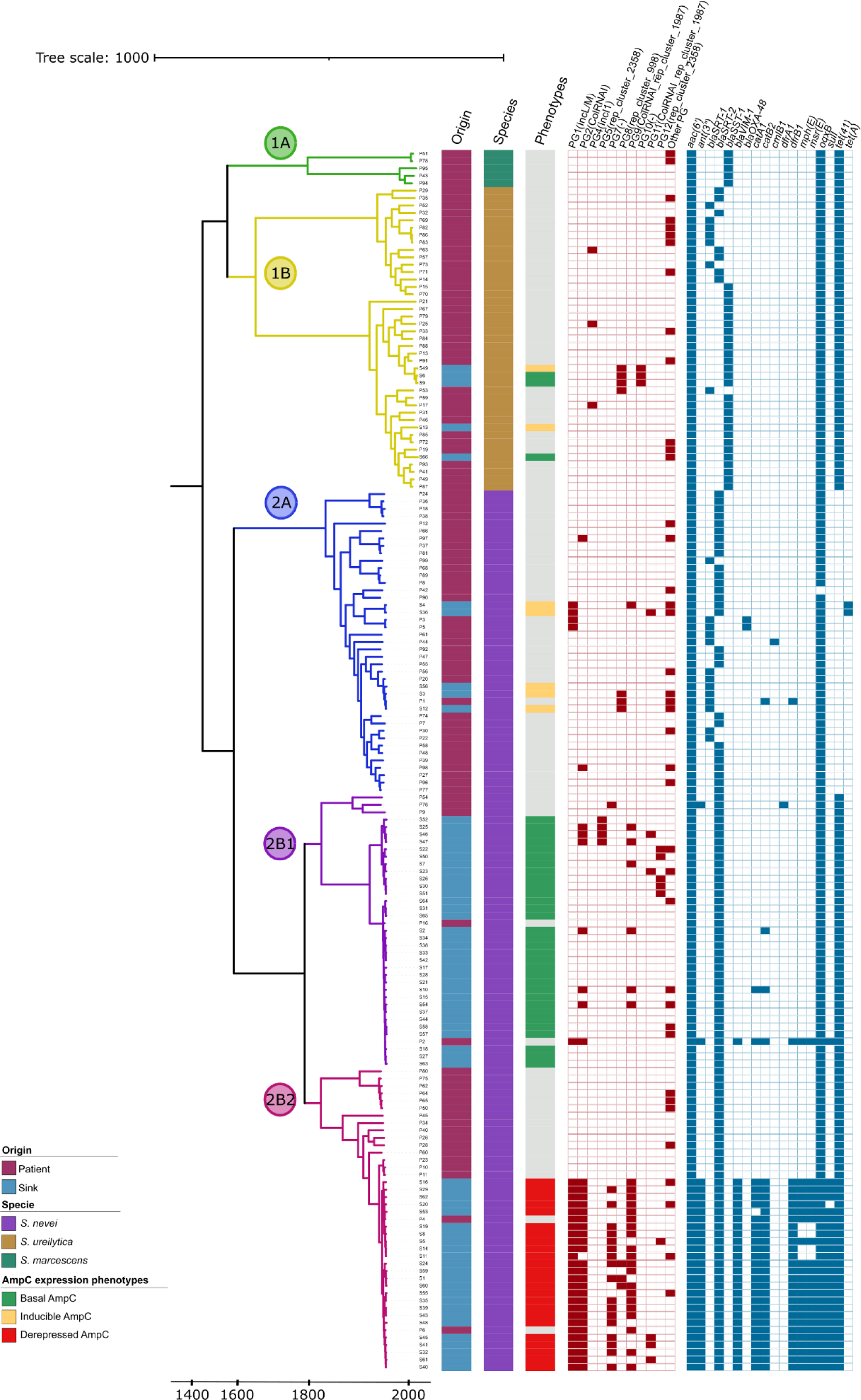
Phylogenetic structure of the 165 isolates of the *Serratia marcescens* complex at Hospital Ramón y Cajal. A maximum likelihood tree built with the 45,720 single nucleotide polymorphisms taken from the 2,583 core genes (appearing in 99% of the genomes). Strains were obtained from sinks and patients (stool, bloodstream, and outbreak-related) from 2003–2020 (see Materials and Methods and Table S1). The main clades and subclades are shaded accordingly (see key). The origin of the sample (patient/sink), the species name according to genome similarity (*S. marcescens, S. ureilytica, S. nevei*), the AmpC phenotype expression (AmpC basal, AmpC inducible, AmpC derepressed, no phenotype tested), the plasmid groups (PG, MOB-Typer), and the antibiotic resistance genes (ResFinder) are represented from left to right and labeled accordingly (see keys).

The single nucleotide polymorphisms (SNPs/Mb) observed in the core genomes of clades and subclades are shown in **Fig. 3**. The “lineage Sn-like” linked to subclade 2A (2003-2020, 0–11,560 SNPs/Mb, median of 9297 SNPs/Mb) includes genomes of highly related environmental isolates from 4/14 ICU sinks and 36 clinical isolates with the AMC^R^+FOX^R^+3GC^S^ phenotype. The “lineage Sn-like” corresponding to subclade 2B was further split into 2B1 and 2B2 (2004-2020, 15834–21784 SNPs/Mb, median of 21338 SNPs/Mb). Subclade 2B1 (0–20107 SNPs/Mb, median of 7359 SNPs/Mb) encompassed genomes corresponding to clonally related environmental isolates from 11/14 sinks carrying an AmpC with basal expression, and 2 clinical isolates collected in 2015 and 2016, one carrying a plasmid harboring *bla*_VIM-1_. Subclade 2B2 comprises the genomes of 18 BSI isolates, 2 clinical outbreak strains, and 23 highly similar environmental isolates exhibiting the AMC^R^+FOX^R^+3GC^R^+4GC^R^±carbapenems^R^ phenotype (VIM-1 producers), which were collected from 7/14 ICU sinks for at least 18 months (0–20107 SNPs/Mb, median of 7359 SNPs/Mb) and persist to date. VIM-1-producing strains from sink and patient outbreaks were highly similar (1–15 SNPs). The “lineage Sm” (subclade 1A, 2003–2016, 0–24943 SNPs/Mb, median of 4528 SNPs/Mb) only include clinical isolates, while the “lineage Su” (subclade 1B, 2003–2015, 0–38265 SNPs/Mb, median of 10038 SNPs/Mb) encompasses 5 environmental isolates from 4/14 ICU sinks with both the AmpC basal and inducible phenotypes, suggesting distinct AmpC expression levels at the strain level. The significance of the values suggests their phylogenomic distance and confirms the long-term persistence of epidemic strains in our dataset. Moreover, the coexistence of *S. nevei* clones exhibiting different PFGE types within subclades 2B1 (basal phenotype) and 2B2 (outbreak VIM-1 producers) indicate the diversity of clonal populations with different genomic conformations. Chromosome alignment revealed the arrangement of a large region (data not shown), which has been observed in comparative genomics studies of *S. marcescens* isolates from different places^2^.

**Fig 3.**
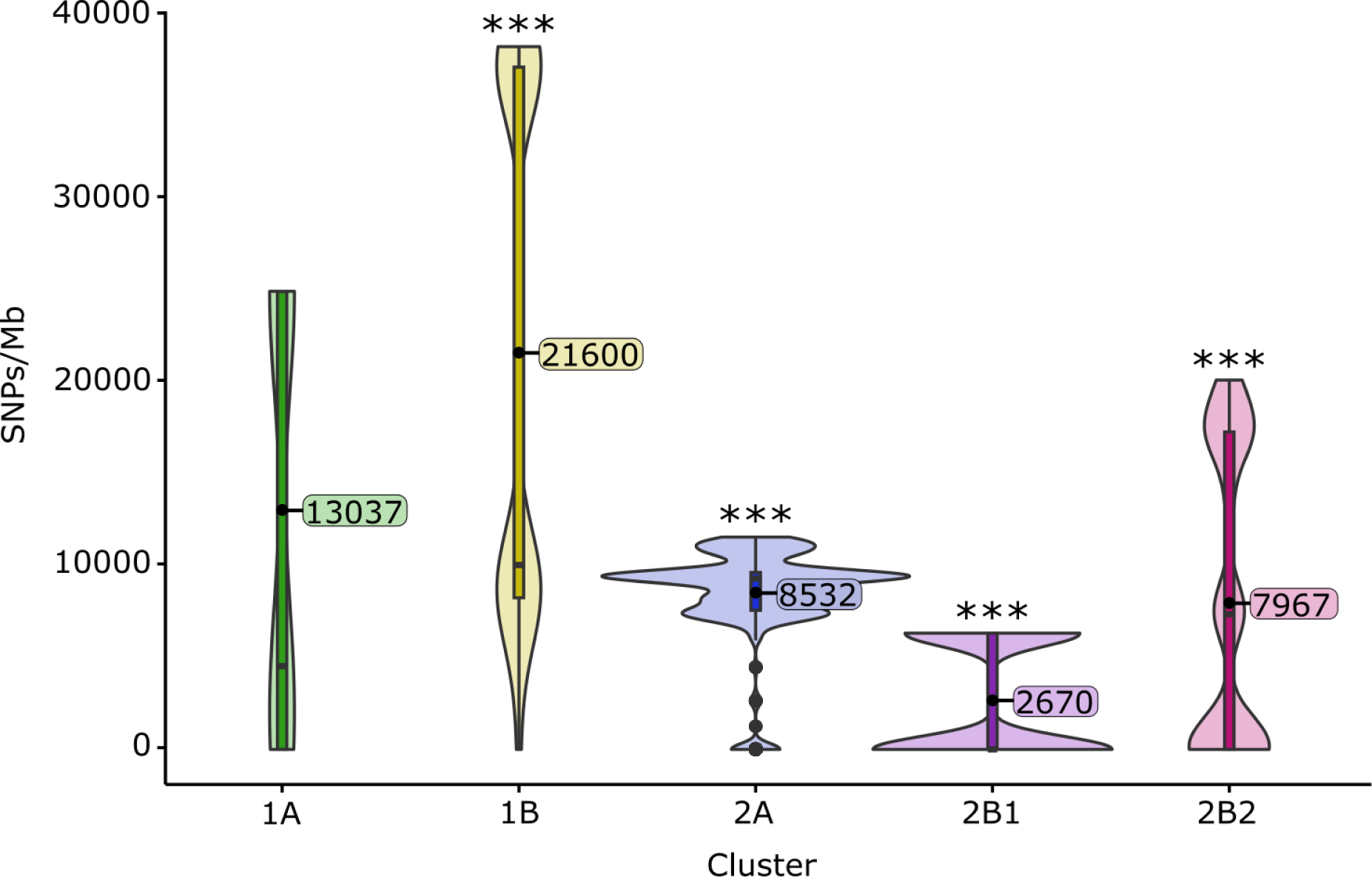
Statistical analysis of the core genome of the *Serratia marcescens* complex. Violin plot showing the single nucleotide polymorphism (SNPs/Megabase [Mb]) distribution among the major clusters of the core *Serratia* genome. SNPs ranged from 35,533 to 45,719 SNPs/Mb between genomes of clades 1 and 2 (42,009 SNPs/Mb median distance), from 0 to 38,432 SNPs/Mb between genomes within clade 1 (33,998 SNPs/Mb median distance), and from 0 to 32,147 SNPs/Mb between genomes within clade 2 (19,303 SNPs/Mb median distance). The significance of the comparisons (Wilcoxon test + Bonferroni p-adjust) was analyzed: 1A-1B (p<0.001), 1A-2B1 (p<0.05), 1B-2A (p<0.001), 1B-2B1 (p<0.001), 1B-2B2 (p<0.001), 2A-2B1 (p<0.001), 2A-2B2 (p<0.001), 2B1-2B2 (p<0.001). Comparisons 1A-2A and 1A-2B2 were not significant (p>0.05).

### The accessory genome

A further analysis of the accessory genomes using the Pangenome Analysis Toolkit (PATO)^21^ (see Materials and Methods for details) was performed to evaluate the presence of adaptive traits in various clades.

We identified 9307 accessory genes (present in less than 80% of the 165 *Serratia* sequences analyzed). **Fig. 4** shows the average number of accessory genes per strain within each subclade (range=931–1159) and the genes shared by strains of the same cluster (range=545–903). A number of genes were significantly enriched within particular subclades (p<0.001) (**Table S2**). Most enriched proteins represent DNA-binding proteins, ATP-binding cassette (ABC) transporters, transcriptional regulators, oxidoreductases, and enzymes involved in lipid metabolism and fimbrial biogenesis. It is of note that the “lineage Sm” (subclade 1A) is enriched in RedY, a homologue to HapK, which is involved in the production of prodigiosin, an emblematic feature of *S. marcescens*^22^. Furthermore, “lineages Sn” corresponding to subclades 2A and 2B2 are enriched in the toxin/antitoxin systems HigB/HigA and HicA/HicB, respectively, while subclade 2B1 only contains antitoxin HigA. These TA systems are enriched in free-living bacteria such as *Pseudomonas* and associated with adaptation to stress responses, biofilm formation and persistence in host and non-host environments^23^.

**Fig 4.**
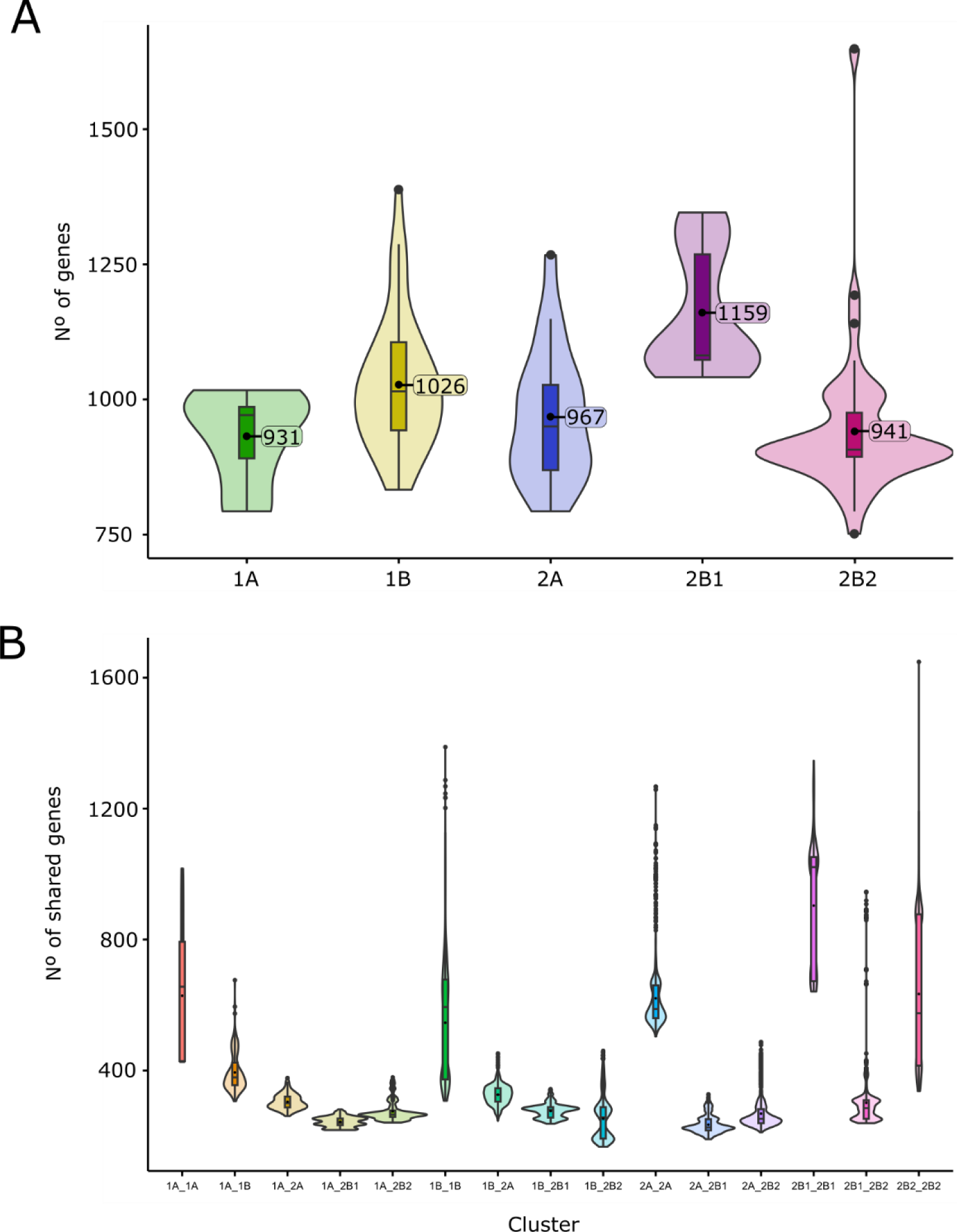
Statistical analysis of the accessory genome of the *Serratia marcescens* complex. **Panel A** represents the number of total shared genes among the clusters (sharedness) calculated for each genome of each cluster and its mean value, employing the Pangenome Analysis Toolkit accent package^21^. **Panel B** shows the number and distribution of accessory genes shared by genomes of the same or different clusters. Each set of genes shared between genomes was evaluated, correcting the model by adjusting the distance between the genomes in order to mitigate the sampling bias. The distribution was modeled with a GLME and Poisson distribution.

A correlation between the structure of the core and the accessory trees was inferred according to the Mantel test (p<0.001). The variables “ward” (the ward in which isolates were recovered) and “origin” (the sample from which strains were isolated) showed a statistically significant association with the phylogeny structures (p<0.001). Interestingly, the variable “ward” showed a higher correlation with the phylogeny (r2=0.2119) than the “origin” (r2=0.1463) of the isolates in the core tree, based on the squared correlation coefficient (r2). In the accessory-based tree, however, the variable “ward” (r2=0.3752) correlated with the sample’s distribution better than the variable “origin” (r2=0.2114).

### The resistome

A comprehensive analysis of the intrinsic and acquired resistome (antibiotics but also heavy metals and biocides) was performed to further explore the contribution of these genes to the adaptability of SMC populations.

*The intrinsic resistome*. All the strains analyzed contained chromosomally located genes associated with resistance to various antibiotic families, such as β-lactams (*bla*_AmpC_), aminoglycosides (*aac(6’)-Ic*), tetracyclines (*tetA*)^11,24,25^, and polymyxins (*pmrA*). All the strains also contained efflux pumps of the resistance-nodulation-division (SdeAB, SdeXY, OqxAB), ABC (SmdAB), and small multidrug resistance (SsmE) families associated with resistance to various antibiotics and/or biocides^26–28^ (**Fig. S2**).

Genotypic and phenotypic variability for certain housekeeping *Serratia* genes was observed among isolates of different SMC clades. Those encoding the cephalosporinase AmpC (and all of their regulatory proteins AmpR, AmpD, AmpE, AmpG) were present in all the genomes; however, mutations in AmpC, AmpR, and AmpD were specific to the different subclades as observed for other species of *Enterobacterales*^13^ (**Table S3**). In addition to the highly conserved AmpC motifs (64SXSK, 150YXN, and 315KTG) and other conserved residues linked to the reference AmpC β-lactamase of *Serratia* SRT/SST, we observed amino acid variations at positions N86K and R91H in the “lineage Sn” linked to subclade 2B, apparently located in the middle of an alpha-helix (H) and a bend or turn^29,30^. A number of mutations (GC>AT) in the intergenic region between the *ampC* and *ampR* genes in these 2B isolates could have increased the number of stop codons affecting the half-life of AmpC and decreased the ability to degrade the antibiotic as suggested^31^. Other changes also observed in AmpC, AmpR and AmpD sequences are shown in **Fig. S3**. The *ant(3’’)*-Ia, previously considered as part of the intrinsic resistome, was only detected in this “lineage Sn” subclade 2B. All isolates but those of the “lineage Sn” subclade 2A contained the allele *tetA*(41).

*The acquired resistome*. The isolates clustered in subclade 2B2 carried a class 1 integron comprising genes that encode resistance to carbapenems (*bla*_VIM-1_), aminoglycosides (*aac(6’)Ib*), chloramphenicol (*catB2*), trimethoprim (*dfrB1*), sulfonamides (*sulI*), and quaternary ammonium salts (*qacEΔa1*) (**Fig. 2**). This gene array (*int*I1*-bla*_VIM-1_*-aac(6’)Ib-dfrB1-aadA1-catB2-qacEΔ1/sul1*) has been detected in a variety of plasmids and species of *Enterobacterales* collected in our hospital for years^16,19^. The IS*26*-*msr(E)*-*mph(E)*-IS*26* transposon conferring resistance to macrolides and triamilides but not lincosamides (and widely spread in the plasmids of *Enterobacterales* and water systems^32^) was detected upstream from the class 1 integron (**Fig. 5C**). Only one other carbapenemase gene, *bla*_OXA-48_, was detected but was confined to clinical outbreak isolates. Of interest was the detection of *pcoABCDRS*, *silABCEFPRS*, and a number of genes of the *arsABCDR* and *mer* operons in the genomes of the subclades 2A and 2B2 (**Fig. S2**). All these metal resistance genes were chromosomally located, in contrast to the plasmid location of contemporary ICU-sink isolates of other *Enterobacterales* species (Guerra-Pinto *et al*, personal communication).

**Fig 5.**
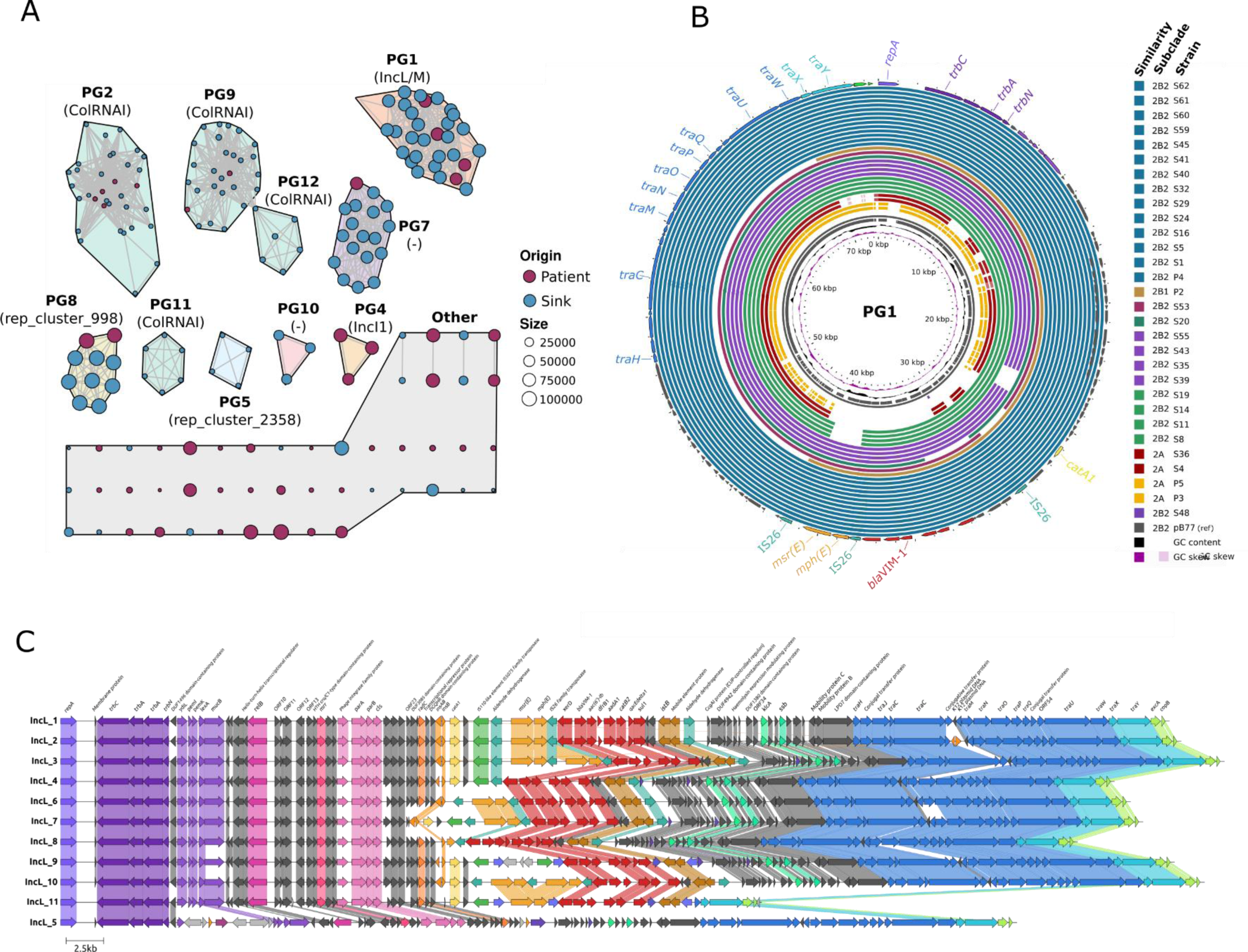
The plasmidome of the *Serratia marcescens* complex. **Panel A:** Plasmid network built with the k-nearest neighbors’ algorithm (K-NNN) using the Pangenome Analysis Toolkit K-NNN function. Each node represents 1 plasmid and is colored according to the source (patient vs. sink), its molecular weight being proportional to the node size (see keys). A plasmid group (PG) was defined using the Louvain algorithm over the network structure, each plasmid being connected with its best hits (Jaccard similarity >0.5, coverage/size difference <50%). PGs were arbitrarily designated with a number; their replicon content as defined by the MOB-typer tool^57^ is shown below the PG label in brackets. **Panel B:** BLAST Atlas of *S. nevei* IncL plasmids in PG1 compared with IncL epidemic plasmids isolated in our institution. BLAST was applied to the coding sequences (CDS) within the reference plasmid from this group, obtained through annotation via Bakta of one of the outbreak plasmid strains (pB77, P6 strain) against sequence regions in the query plasmids. The circularized map of the plasmids was rendered with Gview (https://server.gview.ca/). The inner slots represent the guanine-cytosine (GC) skew, GC content, and CDS regions of the reference. The colored slots represent plasmid data from PG1 strains mostly from subclades 2B2, 2A and 2B1. **Panel C:** Pairwise alignment of all genes using clinker^60^ with GenBank files (.gbff) among 11 variants of the IncL plasmid sequences obtained through MinION technologies and annotated with Bakta. A 100% identity is used for cluster similarity.

### The plasmidome

We identified 190 plasmids among the 165 fully sequenced genomes, of which 132 were clustered in 25 plasmid groups (PGs), and 40 were singletons (**Fig. 5A)**. The predominant PGs in sinks corresponded to known plasmid families L/M (PG1), and Col (PG2, PG9, PG11, PG12) or were not typed by the MOB-suite method (PG7, PG8, PG10). **Tables S4 and S5** shows the features of the 190 plasmids.

Of special interest are PG1 and PG8, which seems to be transferable between sink SMC populations. The PG1 comprises plasmids of high epidemiological value belonging to the L and M families^33,34^. It is overrepresented by a predominant and highly conserved IncL plasmid carrying the class 1 integron mentioned above (**Fig. 5B**) which was highly similar to pB77-CPsm recovered from *Serratia* outbreak isolates since at least 2017^16^. Besides clinical and sink isolates of the “lineages Sn-like” (subclades 2A, 2B1, 2B2), this plasmid was also detected in isolates of *Klebsiella pneumoniae, Klebsiella variicola, Citrobacter cronae,* and *Enterobacter roggenkampii* collected for at least 5 years (2017-2022) from patients^19^, and sinks (Guerra-Pinto *et al*, personal communication). These ∼70 kb pB77-CPsm variants differ in 0–8 SNPs, and/or small indels, rearrangements, or duplications. They were also highly similar to the pOXA-48 circulating in our hospital (99.9% identity, 91% coverage)^17^ encoding *bla*_OXA-48_ instead of the *bla*_VIM-1_ integron (**Fig. 5C**). It is of note that a IncL plasmid variant (∼46 kb) from a clinical isolate, lacking a 27125bp region encoding genes of the conjugation module and categorized as a singleton (“other PGs”), was highly similar to the previous one. PG1 also grouped a ∼81 kb IncM1 plasmid (53.0% GC content) carrying tetracycline resistance genes (*tetA, tetR*) in sink isolates similar to a plasmid previously described in *E. coli*^35^.

The PG8 comprises highly similar ∼107kb plasmids from sink and patient isolates of the “lineage Su” (subclade 1B) and “lineages Sn-like” (subclades 2A, 2B1, 2B2). They were highly similar to p87710 (96.8% identity, 60.0% coverage), a ∼87kb plasmid detected in *S. marcescens* from an oligotrophic pond (GenBank: CP063230.1) but also contained a cluster associated with fimbrial biogenesis (**Fig. S4**). Finally, the PG7 encompasses plasmids of variable size found in our sink and patient isolates and highly similar to pE28_003, a 67kb plasmid isolated in clinical isolate of *S. marcescens* from Australia (GenBank: CP042515) (**Fig. S5**). A variety of Col plasmids (PG2, PG3, PG11, PG12b) were also present in sink and patient isolates from lineages Sn-like (subclades 2A, 2B1, 2B2). Plasmids associated with replicons of the incompatibility group I1 were categorized as PG4 and singletons but they were only detected in clinical isolates.

## DISCUSSION

This work reveals new features about the ecology of *S. marcescens* with implications for controlling and treating *Serratia* infections and mitigating antimicrobial resistance in the healthcare network.

First, the diversity of *S. marcescens* populations as part of the hospital environment’s microbial communities. The study confirms that *S. marcescens, S. ureilytica, S. nematodiphila*, and *S. entomophila* belong to the SMC^9^ and adds *S. nevei* to this complex. The predominance of *S. nevei* probably misidentified in the past as *S. marcescens*^16^ contrasts with the available literature that considers *S. marcescens* the most common “species” in human-associated settings. Since its description in 2020, *S. nevei* has been recovered only from fresh produce in Germany^8^ and from wastewater plants in the US^36^. Other species, such as *S. ureilytica* and *S. nematodiphila*, have rarely if ever associated with humans to date^1^. Specific adaptive traits in core and accessory genomes found for each SMC clade would mirror their diversification and adaptation to disparate conditions and might explain the survival and coexistence of populations in a given environment. Second, the genotypic and phenotypic variability of the AmpC β-lactamase within the SMC. Basal expression of AmpC β-lactamase results in natural full susceptibility to aminopenicillins and cephalosporins, which is rare in species of the ESCPM group and has not been reported in *S. marcescens*^12,13,37^. The association of this basal phenotype with the predominant SMC clade 2B is of clinical importance due to the low probability of AmpC β-lactamase derepression upon therapy^38^. This finding agrees with recent clinical studies that reported low rates of AmpC induction in hospitalized patients infected by *Serratia* and reinforces the need to accurately identify them and test the antibiotic susceptibility prior to treatment implementation^38^. The results would also suggest that AmpC and/or the proteins of its induction cascade would have a role in the adaptation to nonhost habitats. Enzymes involved in peptidoglycan cycling, such as PcgL and DdpX, have been involved in survival outside humans^39^. In *P. aeruginosa* and *Stenotrophomonas maltophilia*, AmpR not only regulates β-lactamases but also proteases, quorum sensing, biofilm formation, and other virulence factors^40–42^. The clade specificity of AmpC β-lactamase and the variability of sequences within clades reflects not only an ancient diversification of the SMC but also a genetic drift, probably as a result of the exposure to multiple selection events in the hospital. Third, hospital water systems enable the “source-sink” dynamics of SMC populations^43^. Hospital sinks, particularly those located in ICUs, are increasingly identified as reservoirs of multidrug-resistant *Pseudomonas* and *Enterobacterales*, with most of the available studies focusing on identifying the origin of outbreaks or the impact of interventions to control bacterial transmission^5,6,44–46^. The absence of long-term analysis of microbial populations with frequent and systematic sampling of patients and sinks during non-outbreak periods precludes the understanding of the transmission pathways of opportunistic pathogens and the ecology of microbial populations in the hospital-built environment. Our study reflects not only the survival and conservation of an epidemic strain producing VIM-1 in sinks long beyond the outbreak but also its coexistence with other SMC lineages and, remarkably, with another persistent and highly susceptible clone of a distinct “*S. nevei*” (subclade 2B1) which was sporadically involved in a polyclonal outbreak of VIM-1 producers in 2017 after acquiring the IncL/pB77-CPsm plasmid^16^. Although the directionality of the transmission is always difficult to demonstrate, the variable occurrence of these long-term persistent “lineages-Sn” in sinks during the study (high recovery during and soon after outbreaks and low recovery when the ICU space was cleared and transformed for other non-healthcare activities) not only suggests that ICU sinks constitute dead-ends of *Serratia* from patients and/or staff but also the ability of *Serratia* to survive in non-host (hospital) environments, an issue only scarcely analyzed for certain *Enterobacterales*^39^. There, they can acquire and act as a reservoir of highly conserved and persistent multidrug-resistant plasmids.

Fourth, the role of *Serratia* in the dynamics of clinically relevant antibiotic-resistant genes and plasmids. The plasmidome of the SMC is still poorly explored, with a few reports describing only the replicon diversity and/or an apparent, however discrete, multidrug-resistant plasmid enrichment among a small number of *S. marcescens* clinical isolates^2,26,47^. The long-term persistence of highly conserved IncL epidemic plasmids carrying *bla*_VIM-1_ or *bla*_OXA-48_ by sink isolates of *Serratia* and other species of the ESPCM group reveals a microbial community that provides population-wide access to broad-host plasmids. This finding contributes to explaining the endemicity of *bla*_VIM-1_ and *bla*_OXA-48_ and plasmid evolution in hospitals far beyond the detailed description of the “units of selection” (clones, plasmid, integrons, transposons) provided by cross-sectional multicentric studies or local reports of polyclonal outbreaks^17,48^. In addition to the IncL plasmids, the distribution of different family of plasmids in sink and clinical isolates (e.g., Col, IncI1 and IncF-like plasmids) mirrors the process by which this species easily acquired, maintained, and enabled plasmid evolution efficiently in various hospital patches, both “source” (permanent) and “sink” (transient) habitats.

In conclusion, the hospital-built environment exemplified by ICU-sinks accounts for diverse and coexisting populations of the SMC able to long survive in host and nonhost environments. Certain SMC clades constitute a unique reservoir of plasmid-carrying genes encoding carbapenemases that can facilitate the endemicity and unpredictable emergence of nosocomial outbreaks involving this species and/or plasmids. From an eco-evolutionary perspective, the findings reflect a “source-sink” dynamics model for SMC lineages and plasmids. The model, used in ecology to understand variations of population growth in heterogeneous habitats such as the hospital ecosystem, has been used to explain antimicrobial resistance by only a few reductionist *in vitro* studies using single plasmids or single clonal backgrounds^43,49^. Here, we demonstrate the relevance of hospital patches (network structure) in the epidemic propagation of clones and plasmids, which is necessary to establish connectivity-dependent infection schemes and to explore the threshold effects where infections would otherwise be unexpected^50^.

## MATERIALS AND METHODS

### Study design and sample collection

The *Serratia* spp. isolates were recovered in the context of a 34-month prospective study in the largest ICU ward (14 rooms, 2 monitor areas) of a tertiary hospital to identify possible environmental reservoirs of opportunistic pathogens. We included 2417 abiotic samples recovered weekly (with a number of discontinuations due to the SARS-CoV-2 pandemic) from sinks (surface, drain, and p-trap, n=1126) and room surfaces (bed rails, ventilator touchscreens, and pots, n=1291). The study comprises 3 periods according to the ward occupancy and the functional uses of the hospital ward. Period A (March 2019-February 2020) and B (July 2020-February 2021) which cover the ICU occupancy before and during the SARS-CoV-2 epidemic, respectively, and period C (March 2021-December 2021, when the patients were cleared out to adapt the ICU ward to other hospital uses and transferred to another floor).

Sink surfaces were sampled with polyurethane sponges placed in a sterile bag impregnated with 10 mL HiCap Neutralizing Broth (EZ-10HC-PUR, EZ Reach^TM^ Sponges, World Bioproducts, Bioing sro, Czech Republic). Sink drainage and p-trap samples were collected with standard sterile swabs and aspiration probes, respectively. The samples were plated onto BD CHROMagar^TM^ Orientation Medium, BD CHROMagar ESBL-Biplate, and mSuperCARBA (Becton Dickinson, Franklin Lakes, USA). The plates were then incubated for 48 h at 37 °C. Colonies of different morphotypes (size, color, and shape) were subcultured onto BD CHROMagar^TM^ Orientation Medium, incubated for 24 h at 37 °C, and further identified with MALDI-TOF MS (MALDI Biotyper, Bruker, Billerica, MA, USA). Stocks of sub-cultured bacterial isolates identified as *Serratia* species by MALDI-TOF-MS (reliability score >2) were stored at −80 °C in 1500 µL Luria Bertani broth + 15% glycerol for further analysis.

In addition to the environmental isolates prospectively recovered in this study, a collection of 99 *Serratia* clinical isolates recovered in our hospital was included for comparative analysis (**Table S1**). There were 94 isolates causing individual episodes of BSI, and 5 VIM-1 and OXA-48 *Serratia* producers involved in hospital outbreaks between 2003 and 2017^16^. Lastly, isolates of *Enterobacterales* carrying *bla*_VIM-1_ plasmids and involved in polyclonal outbreaks^19^ or collected in contemporary studies related to the hospital-built environment were also included (Guerra-Pinto *et al,* personal communication).

### Antibiotic susceptibility

After performing susceptibility testing by disk-diffusion against 14 antibiotics for all 1417 *Serratia* isolates, following CLSI guidelines^51^, we interpreted the phenotypic results by employing criteria from the European Committee on Antimicrobial Susceptibility Testing (https://www.eucast.org/mic_distributions_and_ecoffs/). They included amoxicillin-clavulanic acid (20/10 µg), ampicillin (10 µg), cefoxitin (30 µg), ceftazidime (10 µg), cefotaxime (5 µg), cefepime (30 µg), aztreonam (30 µg), temocillin (30 µg), meropenem (10 µg), ciprofloxacin (5 µg), chloramphenicol (30 µg), gentamicin (10 µg), tobramycin (10 µg), and sulfamethoxazole-trimethoprim (25 µg). The double-disk synergy test (amoxicillin/clavulanic acid and cefotaxime-ceftazidime-cefepime-aztreonam) was performed to screen for ESBL production. Carbapenemase production was phenotypically confirmed with the analysis of a β-lactam phenotype. AmpC β-lactamase induction was screened using cefoxitin and imipenem (medium and strong inducers, respectively) over aminopenicillins (ampicillin), 2GC (cefuroxime), and 3GC (cefotaxime and ceftazidime) as indicators^51^. The presence of carbapenemase genes (*bla*_VIM-1_, *bla*_OXA-48_, *bla*_KPC_, *bla*_NDM,_ and *bla*_GES_) was tested with a multiplex polymerase chain reaction assay^52^.

### Clonal relationship

A clonal relationship between isolates was preliminarily established by PFGE, using 3000 U of XbaI (Takara, Japan) following the PulseNet website protocols (https://pulsenetinternational.org/protocols/pfge/). Comparison of XbaI-digested genomic patterns revealed various pulsotypes (>4 bands) from which we selected a set of isolates for further typing by whole genome sequencing according to spatiotemporal distribution, pulsotype (PT), and antibiotic susceptibility.

### Genome sequencing

All isolates were sequenced by Illumina (we sequenced 66 “de novo” sink isolates plus 93 from the strain collections of the microbiology department). We further closed the sequence of 23 isolates using long-read sequencing (Oxford Nanopore) to accurately determine chromosome and plasmid synteny. We cultured isolates from frozen stocks (−80 °C) on LB agar overnight at 37 °C and performed DNA extraction from 2 mL of liquid subcultures. For short-read sequencing, we used the Chemagic DNA Bacterial External Lysis Kit (PerkinElmer, USA) following manufacturer recommendations. We measured DNA quality and concentration in a NanoDrop 2000 Spectrophotometer (Thermo Scientific, Waltham, MA, USA) and Qubit® 2.0 Fluorometer (Life Technologies, Waltham, Massachusetts, USA) and assessed fragment length with the TapeStation 2200 (Agilent,Waldbronn, Germany). We performed fragmentation of 4 µg DNA in a 46-µL elution volume in Covaris G-tubes by centrifuging at 4200–5000 rpm (Eppendorf 5424 centrifuge) for 90 s to achieve fragment sizes of ∼20 kb. We prepared DNA libraries employing the Nextera XT library preparation kit and the Nextera XT v2 index kit (Illumina, San Diego, CA, USA). The library was sequenced on a HiSeq4000 (Illumina) using the reagent kit v2 (Oxford Genomics Center, Oxford, UK) to generate 250-bp paired-end reads.

For long-read sequencing, we extracted DNA with the MagnaPure 96 System (Roche, Basilea, Switzerland) and quantified it with a Qubit^®^ 2.0 Fluorometer. We used a minimum concentration of 20 ng/µL for the library preparation, employing the Rapid Barcoding kit 96 (Oxford Nanopore Technologies, Oxford, UK), following the manufacturer’s instructions. We loaded the libraries onto flow cell versions FLO-MIN106 R9.4 MinION and sequenced them for 72 h. Base calling was performed in real-time with Guppy integrated in MinKNOW (Oxford Nanopore). Adapter sequences were removed with qcat (Oxford Nanopore), and fastq reads were filtered and quality measured with NanoFilt (>10,000 bases) and Filtlong v0.2.039 (500Mbp best reads) before the assembly. Hybrid assemblies were created with Unicycler v0.4.7^53^ and checked by visualization with Bandage v0.8.1.

### Genomic analysis

The 165 sequenced genomes were first annotated with Bakta 1.7.0^54^. We then defined the core and accessory genomes with the Pangenome Analysis Toolkit R-package, which enabled us to analyze the population structure, annotate adaptive features, and create gene networks^21^. The function “classifier” was employed to search for the closest reference species in the NCBI. Further average nucleotide identity (ANI) analysis led to establishing the taxonomic and phylogenetic relationship between various *Serratia* subclades. A distance matrix was created with ANI distances calculated using PATO. A dendrogram was generated with the “tree_dist” command in PATO. Clades were defined based on minimum within-node pairwise ANI scores using the unsupervised clustering MClust^55^. The size of the pangenome, core, and accessory genome was calculated with the function “core_plots” of PATO.

### Core genome

We identified 2583 core genes (blastP identity cutoff 90%, the core genes being defined as those present in >90% of the isolates) and analyzed through a pseudo-multisequence alignment with FastreeMP^55^ midpoint root to build a maximum likelihood phylogenetic tree. The tree was displayed and annotated using ggtree version 2.2.4 in the R package. We employed iTOL^56^ to edit and generate metadata linked to the tree.

### Accessory genome

We determined the accessory genome employing PATO with default parameters (80% identity and 80% coverage), defined as the set of genes detected in less than 80% of the genomes^21^. We further analyzed the accessory gene enriched in each subclade, using the PATO function “accnet_enrichment_analysis”, which performs a multi-hyperparametric test to find overrepresented genes in a cluster compared with the general population (i.e., the genome data set). The accessory network was illustrated with Gephi for rearranging and uploading the layout in R to plot the network. We performed a statistical test of accessory genes shared within each CG or within each subclade, modeling the distribution of gene pair sharedness between genomes of each clade/subclade, which was further corrected according to the similarity between genomes. For the statistical test of the number of accessory genes per genome, we employed a nonpaired Student’s t-test (stats R package).

### Plasmid analysis

Plasmids were reconstructed from FASTA files, typed using the MOB-recon and MOB-typer modules of the MOB-suite software^57^, and annotated with Bakta^54^. We then created a distance matrix using gene-by-gene presence-absences using the “accent” function of PATO with a default similarity parameter to 70%. That distance is performed using Jaccard similarity. We then created a k-nearest neighbor network (K-NNN) to allow reciprocal connections with the thresholds of 10 neighbors and 0.5 Jaccard distance. This implies that any plasmid is linked to its 10 most similar plasmids provided they share more than 0.5 Jaccard similarity. Plasmids were clustered from the K-NNN using mclust^55^.

The plasmid network was built with Cytoscape, imported to R with the tidygraph R package, and set with the Louvain cluster algorithm (igraph R package). The plasmid typing information (predicted mobility, replicases, and relaxases) was added to the network. Plasmid distribution over phylogenetic trees was built with ggtree R software. All data manipulation and visualization were performed with the tidyverse R metapackage.

Hybrid assemblies of plasmid sequences were submitted to Plasmidfinder^33^ and Resfinder^58^ for replicase and antibiotic gene identification, and Biocides and Metals from Bacmet with Blast against the BacMet database^59^. Gview (https://server.gview.ca) was employed for generating the circular map and clinker^60^ for the linear visualization and gene cluster comparison.

### Statistics

The Mantel test implemented in R was employed to compare the structure of the core and accessory trees by determining the correlation between the distance matrices. To evaluate the association of the metadata variables (origin, collection place, and year) with the core and accessory tree, we employed the “envfit” function in the vegan R package. This function calculates the multiple regression of environmental variables with the ordination axes and estimates the significance with a permutation test. The p-values were adjusted based on the Bonferroni correction. The goodness-of-fit statistic is the squared correlation coefficient (r2), which measures the correlation of the variable with the ordination.

### Accession numbers

All sequences were deposited in the European Bioinformatics Institute database under bioproject accession number PRJNA1023224 (accessible from January 1^st^, 2024). WGS data from outbreak clinical strains were previously submitted^16^ under the accession numbers JAAAMA000000000, JAAAMB000000000, JAAAMC000000000, JAAAMD000000000, and JAAAME000000000. Closed plasmids were fully sequenced and analyzed during this work under the same bioproject.

## ACKNOWLEDGMENTS

We would like to thank Astrid Rasmussen for the hybrid assembly of the genomes and plasmids. During the implementation of this study, SAG and SSC were supported by the Youth Employment Operational Program of the Regional Government of Madrid (PEJD-2019-PRE/BMD-15530 and PEJ-2020-TL/BMD-18185, respectively). MDFdB and NGP were supported by the Instituto de Salud Carlos III (pFIS F19/00366 and PI18/1942, respectively).

## AUTHORS’ CONTRIBUTION

SAG: wet lab (identification and typing of *Serratia* environmental isolates), Minion sequencing, genomic and plasmid analysis, and manuscript writing; MDFdB: bioinformatics analysis of core and accessory genomes, plasmidome, revision of the manuscript; NGP and SSC: sampling and typing of sink isolates; AEPC: statistical analysis and revision of the manuscript; RC: provision of isolates from strain collections of the Microbiology Department and valuable information from the clinical microbiology lab, discussion of the results, revision of the manuscript; BPV: information regarding clinical BSI isolates; SR: discussion of the work and revision of the manuscript; CS and RdP: accessibility to the ICU, valuable information about ICU metrics; VFL: direction and supervision of all bioinformatics analysis and manuscript revision; HH: characterization of IncL plasmids by hybrid sequencing approaches; FB: study design, data analysis and revision and edition of the manuscript. TMC: study design, data analysis, manuscript writing. All the authors have read and approved the final version of this document.

