## Supplementary Material for "Long-term dynamics of the “*Serratia marcescens* complex” in the hospital-built environment"

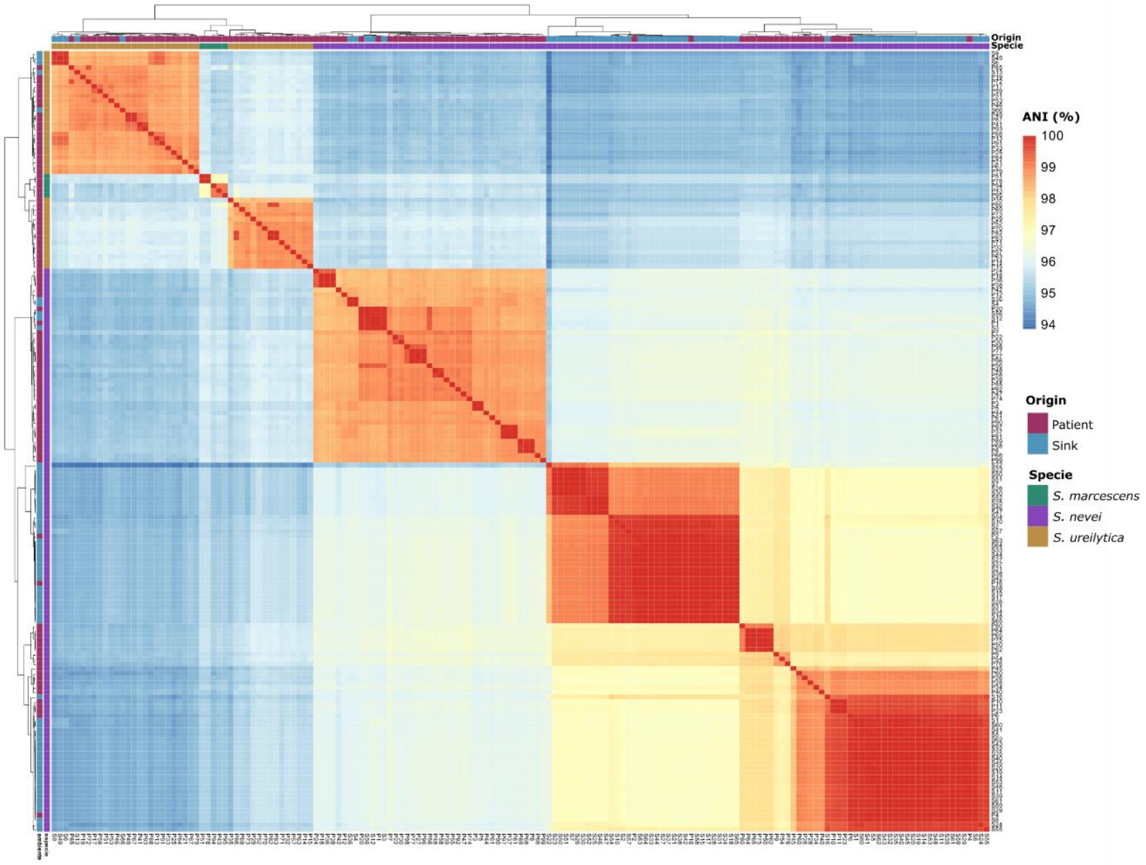


**Fig. S1.** Average nucleotide identity (ANI) between *Serratia* isolates encompassed in clades 1A (*S. marcescens*), 1B (*S. ureilytica*), and 2 (*S. nevei*), represented by a heatmap. The color bar represents the ANI value, whereas the lines bordering the figure represent source and species (from outside to inside). Heatmap of ANI from the core genome of *Serratia*.

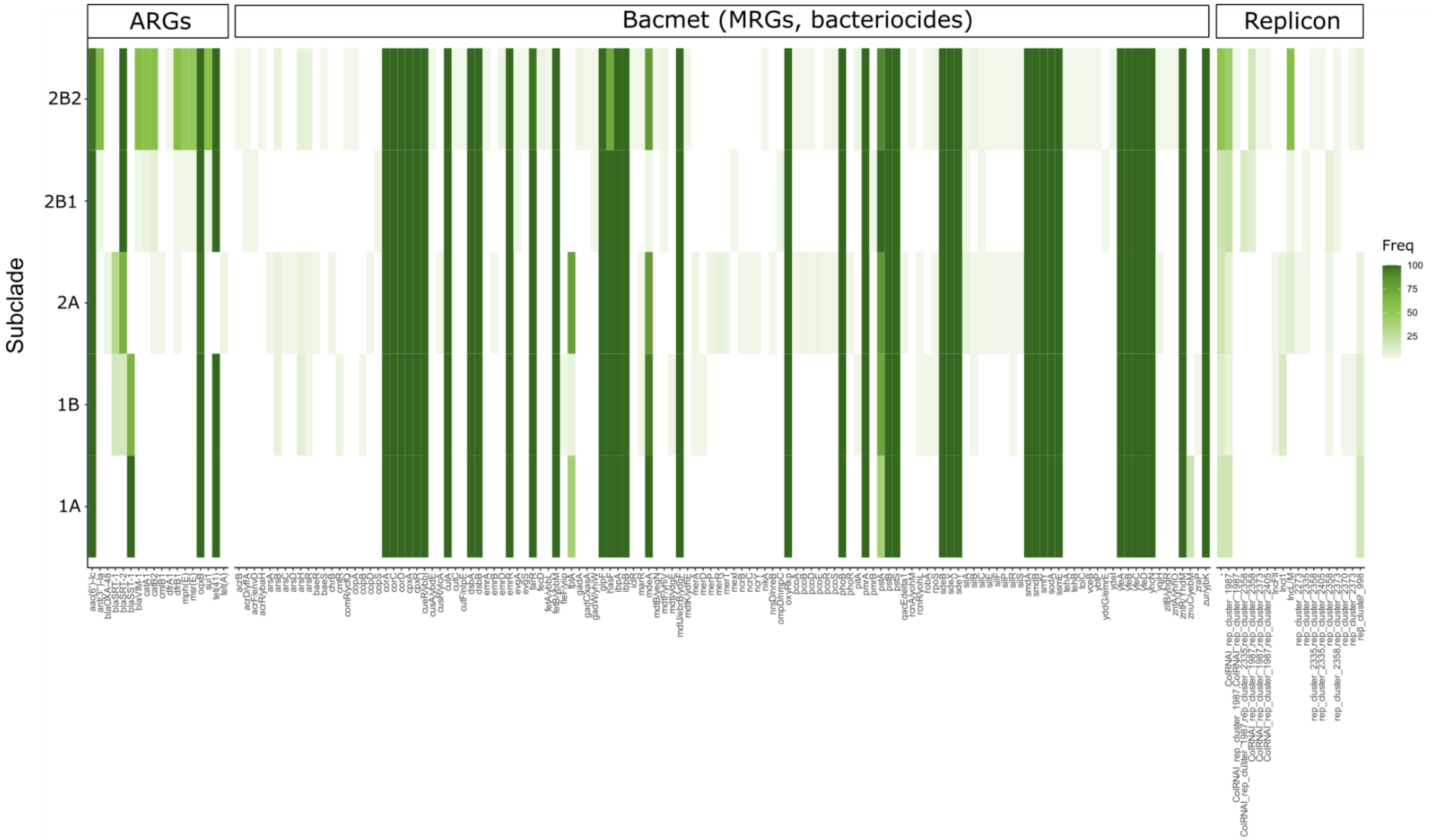


**Fig. S2**. Resistome, metalome and plasmidome of SMC isolates. The abscissa shows the isolates grouped in the "lineages Sm "(subclade 1A), "lineage Su" (subcluster 1B), and "lineages Sn-like" (subclades 2A, 2B1,2B2). The ordinate axis represents ARGs (antibiotic resistant genes) and MRGs (metal resistance genes), and plasmid replicons. They were identified after interrogating the ResFinder (https://cge.food.dtu.dk/services/ResFinder/), BACMET (http://bacmet.biomedicine.gu.se/advanced_search.pl) databases, and MOBsuite databases.

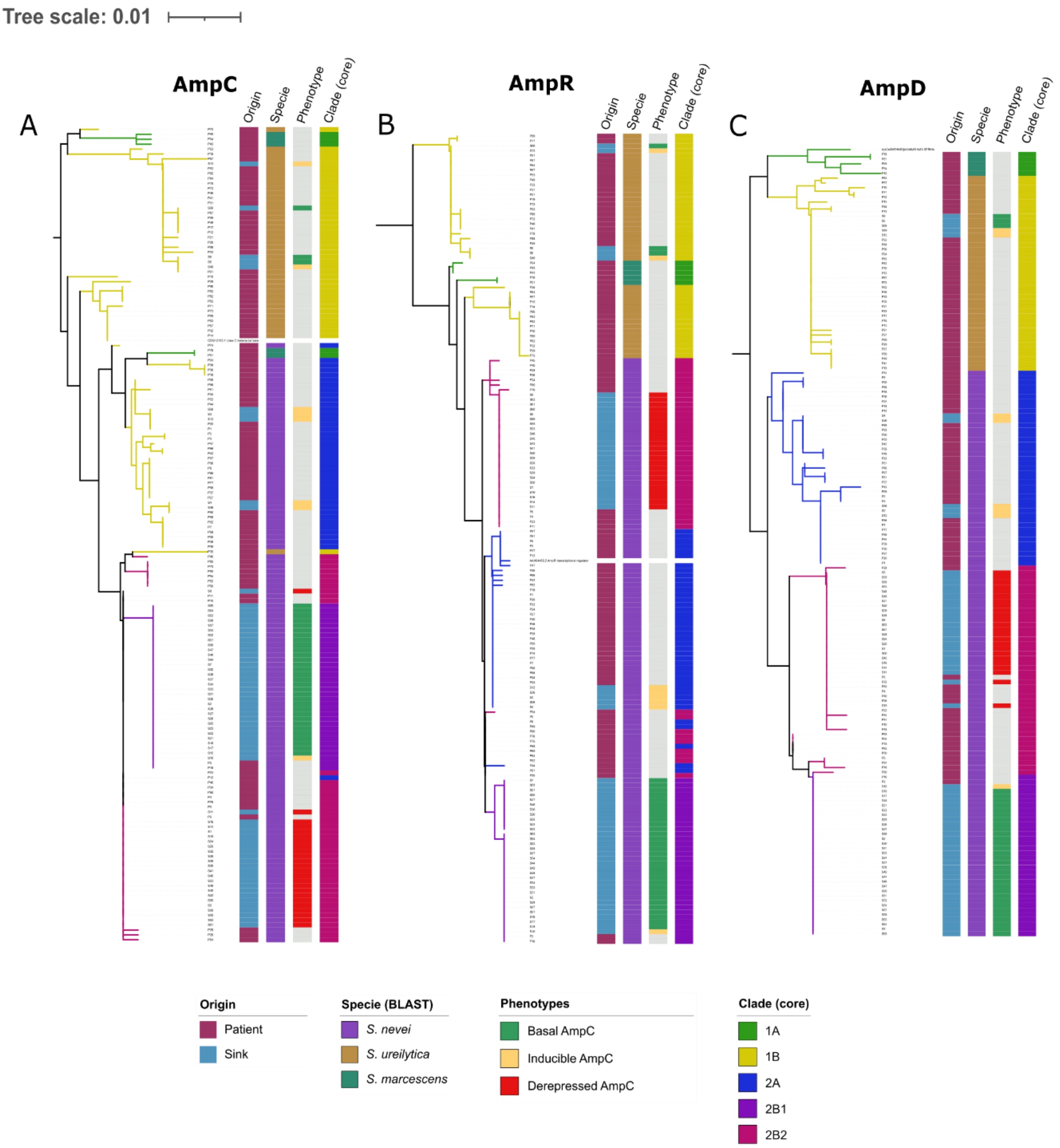


**Fig. S3.** Phylogenetic tree of a multiple sequence alignment of the AmpC, AmpR, and AmpD protein sequences of *Serratia* spp. (panels A, B, C, respectively) generated by ClustalW (https://www.ebi.ac.uk/Tools/msa/clustalo/). The epidemiological features of the isolates (β-lactam phenotype, sample origin, *Serratia* lineages and clades/subclades inferred from the core genome phylogenetic tree) are represented in bars (see keys).

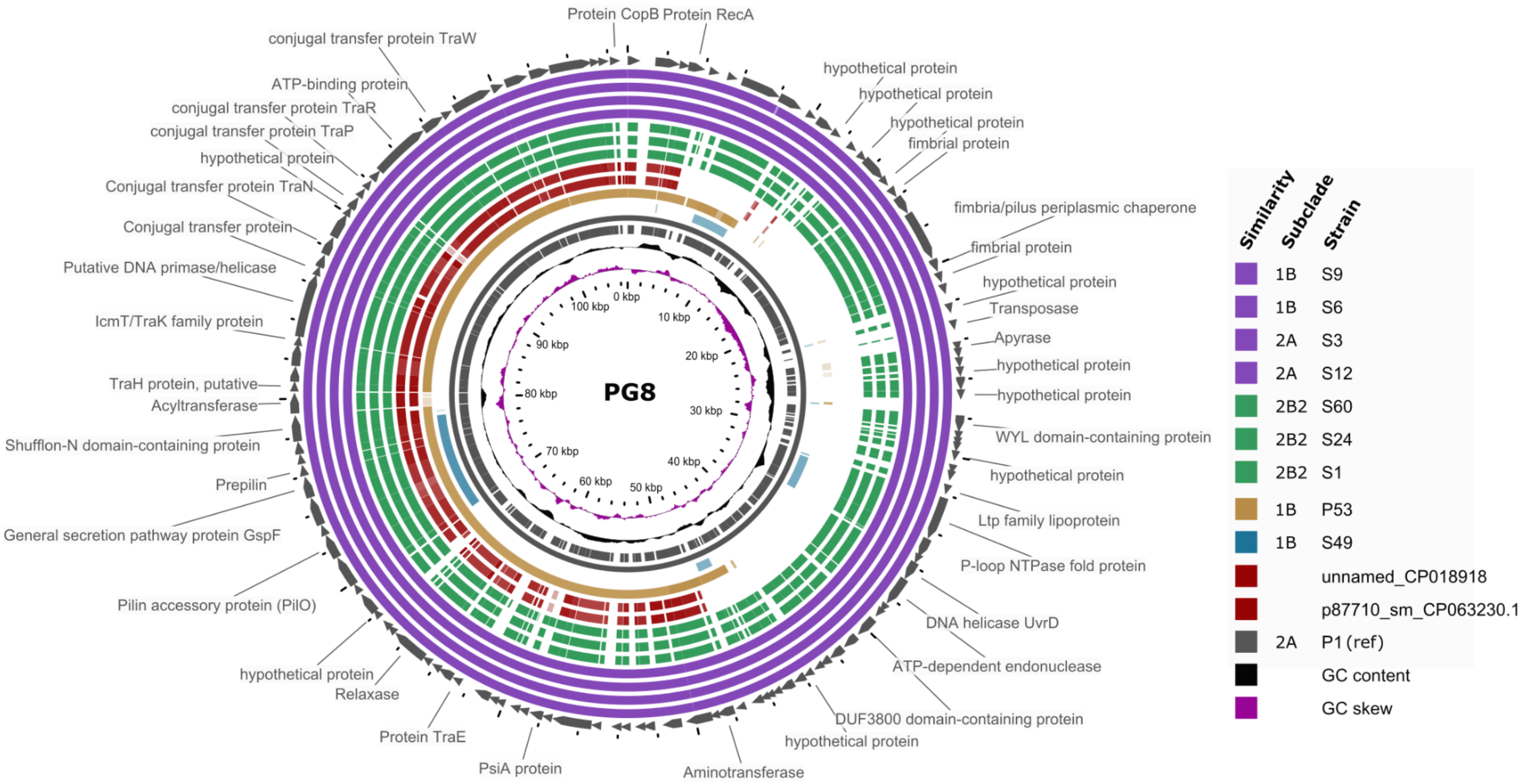


**Fig. S4**. Basic Local Alignment Search Tool (BLAST) Atlas of *Serratia* IncF-like plasmids in PG8. BLAST was applied to the coding sequences (CDS) within the reference plasmid from this group, obtained through annotation via Bakta (S60_plasmid2) against sequence regions in the query plasmids. The circularized plasmid map was rendered with Gview (https://server.gview.ca/). The inner slots represent the guanine-cytosine (GC) skew, GC content, and CDS regions of the reference.

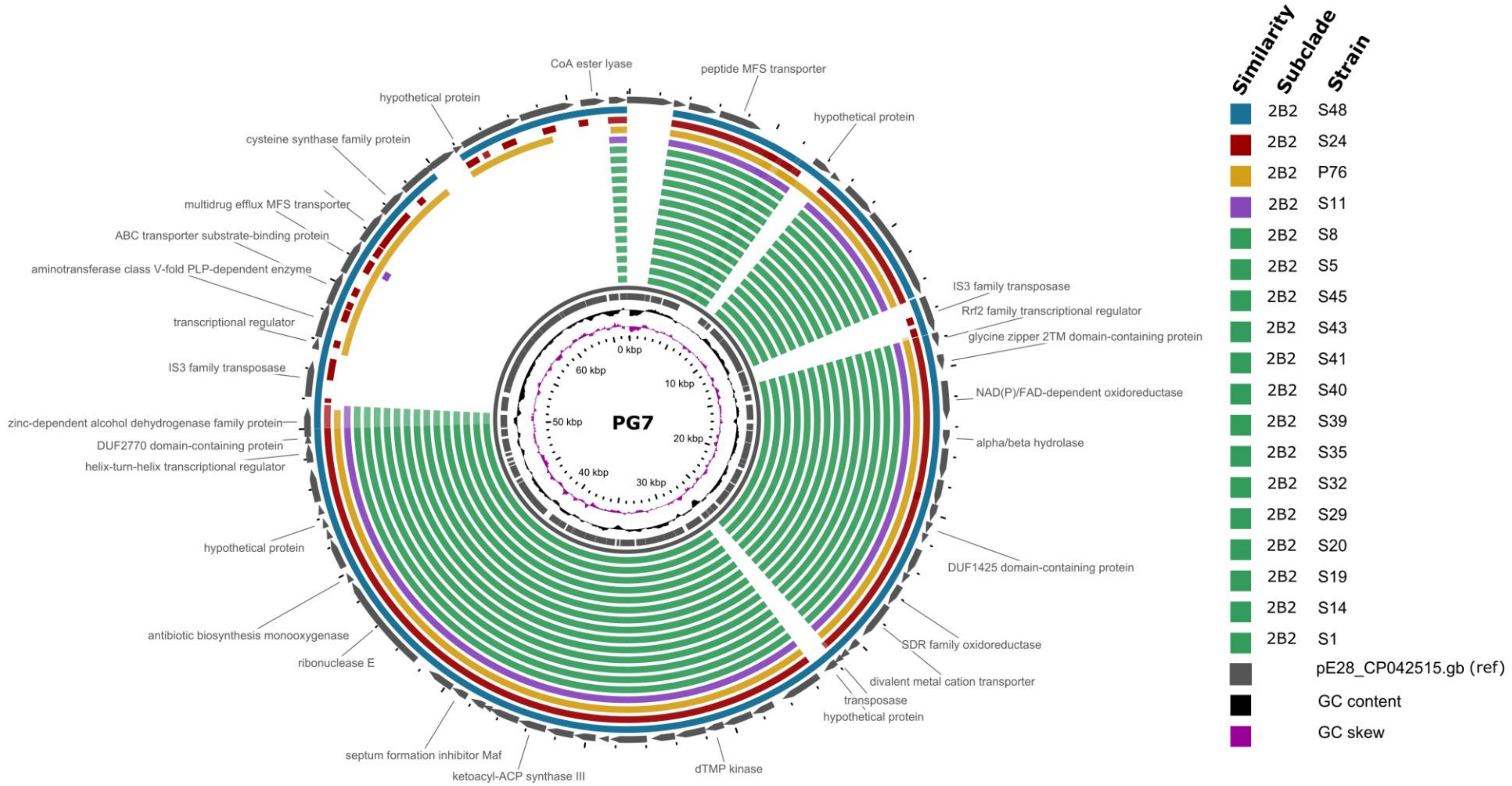


**Fig. S5.** Basic Local Alignment Search Tool (BLAST) Atlas of *S. nevei* plasmids in PG7. BLAST was performed to the coding sequences (CDS) within the reference plasmid from this group (pE28, GenBank: CP0425, most similar plasmid obtained through MOBTyper) against sequence regions in the query plasmids. The circularized map of the plasmids was rendered with Gview (https://server.gview.ca/). The inner slots represent the guanine-cytosine (GC) skew, GC content, and CDS regions of the reference.

**Table S4**. Plasmid Group (PG) dataset

| | | | | | | | Origin | | Subclades | | | | |
|---|---|---|---|---|---|---|---|---|---|---|---|---|---|
| **Plasmid Group (PG)** | **Rep-type*** | **PG size (n)** | **Size (kb)** | **Relaxase_ Type (n)** | **Mash_ ID** | **Mash_ identification** | **C** | **S** | **1A** | **1B** | **2A** | **2B1** | **2B2** |
| **PG1** | IncL/M | 31 | 44.0-74.6 | MOBP(30), -(1) | CP023251, CP023419, KX230795 | Kp(29), Ecc (2) | 5 | 26 | 0 | 0 | 2 | 1 | 28 |
| **PG8** | rep(IncF-like) | 10 | 96.3-107.1 | MOBP(10) | CP018916 | Sm(8) | 2 | 8 | 0 | 4 | 3 | 0 | 3 |
| **PG7** | - | 18 | 62.2-64.4 | -(18) | CP042515 | Sm(18) | 1 | 17 | 0 | 0 | 0 | 0 | 18 |
| **PG10** | - | 3 | 37.9 | MOBP(3) | CP018921 | Sm(3) | 0 | 3 | 0 | 3 | 0 | 0 | 0 |
| **PG4** | IncI1** | 3 | 55.2-80.7 | MOBF(3) | LT575492 | Sm(3) | 3 | 0 | 0 | 3 | 0 | 0 | 0 |
| **PG5** | rep_2358 | 4 | 3.6 | MOBP(4) | CP034043 | Kpp(4) | 0 | 4 | 0 | 0 | 0 | 4 | 0 |
| **PG2** | ColRNAI | 34 | 1.5-2.4 | -(34) | CP041737 | Ehs(34) | 5 | 29 | 0 | 0 | 2 | 7 | 25 |
| **PG9** | ColRNAI | 27 | 2.8-5.9 | MOBP(27) | CP016867, CP024503, CP024519 | Kp(2), Set(25) | 2 | 25 | 0 | 0 | 1 | 6 | 20 |
| **PG11** | ColRNAI | 6 | 3.3 | MOBP, MOBP(6) | CP016867 | Set(6) | 0 | 6 | 0 | 0 | 1 | 2 | 3 |
| **PG12** | ColRNAI | 6 | 3.3-5.8 | MOBP(6) | CP016867 | Set(6) | 0 | 6 | 0 | 0 | 0 | 5 | 1 |
| **Other** | multiple | 48 | 1.1-107.1 | multiple | multiple | multiple | 31 | 17 | 3 | 14 | 16 | 7 | 8 |

**Abbreviations:** Kp (*Klebsiella pneumoniae*), Kpp (*Klebsiella pneumoniae* subsp. p*neumoniae),* Ecc (*Enterobacter cloacae* subsp. *Cloacae),* Ehs (*Enterobacter hormaechei* subsp. *Steigerwaltii),* Sm (*Serratia marcescens),* Set (*Salmonella enterica* subsp. *enterica* serovar *Typhimurium* str. CDC 2010K-1587), C (clinical isolates), S (sink isolates). **Notes**: *Rep-type according to MOBTyper[57]; **IncI plasmids were also identified in small PGs besides PG4. They include PG18 and PG28 (subclade 1B), PG6 (subclade 2A), and PG3 (subclades 1B and 2A), all corresponding to clinical isolates.

Detailed description of each plasmid and plasmid groups appears in Table S5
